# Validation of Body Condition Scoring as a Screening Test for Low Body Condition and Obesity in Common Marmosets (*Callithrix jacchus*)

**DOI:** 10.1101/2025.03.18.643796

**Authors:** Juan Pablo Arroyo, Addaline Alvarez, Lori Alvarez, Alexana J. Hickmott, Aaryn C. Mustoe, Kathy Brasky, Kelly R. Reveles, Benjamin J. Ridenhour, Katherine R. Amato, Michael L. Power, Corinna N. Ross

**Affiliations:** Southwest National Primate Research Center, Texas Biomedical Institute, San Antonio, TX, USA; College of Pharmacy, The University of Texas at Austin, Austin, TX, USA; Graduate School of Biomedical Sciences, University of Texas Health, San Antonio, San Antonio, TX, USA; Department of Mathematics and Statistical Science, University of Idaho, Moscow, ID, USA; Department of Anthropology, Northwestern University, Evanston, IL, USA; Center for Species Survival, Smithsonian’s National Zoo and Conservation Biology Institute, Washington, DC, USA

**Keywords:** body composition, quantitative magnetic resonance, obesity, low body condition, body condition scoring, screening test

## Abstract

Assessing body weight is common practice for monitoring health in common marmosets (*Callithrix jacchus*). Body composition analysis via quantitative magnetic resonance (QMR) is a more in-depth assessment allowing measurements of lean and fat mass, but it is expensive and remains unavailable to most. Alternatively, body condition scoring (BCS) is an instrument-free method for visually inspecting and palpating lean and fat tissue. Animals are rated for lean and fat mass abundance, using an ordinal scale with species-specific descriptions as reference. However, modified BCS systems developed for other species are being used, because no BCS system has been fully validated for marmosets. The accuracy of BCS in identifying marmosets with poor body condition or obesity remains unknown. We assessed an adapted BCS for marmosets (n=68, 2–16 years). Objectives were to 1) determine whether BCS predicts body weight and body composition, and 2) evaluate the performance of BCS as a screening test for low body condition and obesity in marmosets, in comparison to QMR body composition analysis. BCS predicted body weight and body composition (*F*(15, 166)=7.51, Wilks’ Λ=0.240, *p*<0.001), and was better at predicting low lean mass and obesity, than at predicting low adiposity. Marmosets with low BCS had higher odds of low lean mass (*B*=3.37, (95% CI, 0.95-5.78), OR=29.0, *p*=0.006). Marmosets with excessively high BCS had higher odds of obesity (*B*=2.72, (95% CI, 1.07-4.38), OR=15.23, *p*=0.001). The accuracy of BCS suggests it can serve as an instrument-free method to screen for low body condition (79.4%-91.2%) and obesity (77.9%) in marmosets.

**Research highlights:**

- We evaluated body condition scoring (BCS) as a screening tool for detecting low body condition and obesity in marmosets by comparing it to diagnoses based on quantitative magnetic resonance, the gold-standard method for body composition analysis.
- BCS was more accurate at detecting low lean mass and obesity than low adiposity, with marmosets having low BCS showing higher odds of low lean mass and those with excessively high BCS having higher odds of obesity.
- Results suggest that BCS can serve as an instrument-free method to screen for low body condition and obesity in marmosets, enabling early detection of health decline and guiding the need for further diagnostic testing and treatment.

## Introduction

*Callithrix jacchus* (common marmosets) are increasingly employed in biomedical research as a model for aging, health and disease (Miller et al., 2016; Ross, Colman, Power, & Tardif, 2020; Ross & Salmon, 2019). Keeping track of body weight has been an essential part of preventive care for marmoset populations, as wasting and obesity usually result in adverse health outcomes. Marmoset wasting syndrome has historically been a problem in captivity associated with weight loss, weakness, diarrhea, anemia, nephritis, and increased mortality (Brack & Rothe, 1981; Cabana, Maguire, Hsu, & Plowman, 2018; Ludlage & Mansfield, 2003; Otovic, Smith, & Hutchinson, 2015; Yamazaki et al., 2020). At the other extreme, the occurrence of obesity (defined as ≥10% body fat) (Power, Ross, Schulkin, & Tardif, 2012) in captive colonies has become a concern, as obese animals are more likely to exhibit dyslipidemia, altered glucose metabolism, hepatic steatosis, atherosclerosis, cardiomyopathies, and stroke (Power, Ross, Schulkin, Ziegler, & Tardif, 2013; Ross et al., 2020; Tardif et al., 2009).

Monitoring body weight and body composition can provide insights about the physical state of a marmoset. Body weight is the measurement of total body mass. Body composition analysis is a more in-depth assessment capable of distinguishing between lean mass and fat mass in an organism. Measurements in body composition analysis include absolute values, as well as proportional values between lean mass, fat mass, and total body weight. In this sense, body composition analysis is a more comprehensive approach as it generates measures that are specific and proportional; while body weight is a single non-specific measurement.

Focusing on body weight without considering body composition, can result in misclassification of individuals, and may become problematic when making comparisons at the individual and population levels. Individuals with the same body weight and body size might fall into different body composition categories (i.e., have different proportions of lean and fat mass) and exhibit a different health status or risk for disease. For instance, assessments relying only on body weight might suggest that an individual with average weight is “normal” or “healthy”. However, with body composition analysis it is possible to identify average-weight individuals that have proportionally high values of fat (i.e., obesity) or low values of lean mass (i.e., muscle wasting). Consequently, using body weight as a proxy for adiposity can result in underestimation or overestimation of fat mass. Body composition analysis is better suited for developing categories, assigning individuals to categories, and making comparisons across populations; as it provides a more nuanced and complete picture of an organism’s body mass and how it may relate to health.

Quantitative magnetic resonance (QMR) has been shown to be a reliable non-invasive method for acquiring precise measurements of body composition parameters in marmosets (Power et al., 2012; Tardif et al., 2009). However, the equipment required for QMR body composition analysis is expensive and may not be readily available to most clinicians and researchers. As a result, body weight and morphometric measurements are more frequently used for assessing marmosets, instead of QMR body composition analysis – potentially leading to inaccurate estimations of adiposity and misclassifications for disease risk.

The term “body condition”, which is related but not synonymous with body composition, is often used to describe the general state of an individual, emphasizing the importance of fat and lean mass abundance. The concept of body condition is based on the premise that the overall physical state or appearance of an organism can be indicative of energetic reserves, health, and ability to cope under physiologically challenging conditions (Schulte-Hostedde, Millar, & Hickling, 2001). It follows that visual examination and palpation can provide insights into an organism’s body condition, thus measures frequently used as proxies for the construct of body condition include body weight, morphometrics, and body condition scoring (Labocha & Hayes, 2012).

Body condition scoring (BCS) is a tool for rapid and instrument-free assessment, widely used in wildlife biology, veterinary medicine, and domestic and exotic animal management (Burkholder, 2000; Joblon et al., 2014; Labocha & Hayes, 2012; Schiffmann, Clauss, Hoby, & Hatt, 2017). BCS systems are subjective semi-quantitative procedures that consist of visually inspecting and palpating fat and lean tissue on distinct anatomical landmarks. Individuals are then rated, using an ordinal scale with species-specific descriptions as reference (Clingerman & Summers, 2005). BCS systems have been studied and validated for use in several non-human primate species (Ghassani et al., 2023; Reamer et al., 2020; Summers, Clingerman, & Yang, 2012; Torfs et al., 2023). However, marmoset-housing institutions have been relying on modified versions of BCS systems that were originally developed for other non-human primate species, as no BCS system has yet been fully validated for marmosets. The availability of a validated BCS system for marmosets may improve the capability for early detection of animals that may be at risk, as well as facilitate methodological standardization across institutions (Goodroe et al., 2021).

The validation of a marmoset BCS system depends upon its accuracy as a screening test (Maxim, Niebo, & Utell, 2014; Trevethan, 2017) to detect both extremes of body condition, low body condition (e.g., wasting) or obesity. Screening tests are evaluated for accuracy by comparing results against a “gold standard” or benchmark test, which is a widely accepted reliable method for diagnosing a specific disease or condition (Maxim et al., 2014; Trevethan, 2017). In this case, QMR body composition analysis represents a non-invasive gold standard method for precise measurements of lean and fat mass in marmosets.

Here we assessed an adapted version of a BCS system for marmosets, suggested by Burns and Wachtman (2019). The objectives of this study were to 1) determine whether this BCS system accurately predicts body weight and body composition parameters in marmosets, and 2) evaluate the performance of BCS as a screening test for low body condition and obesity in marmosets, by comparison to QMR body composition analysis.

## Methods

### Subjects

In this study we examined a sample of 68 common marmosets (*Callithrix jacchus*), composed of 35 females and 33 males between 2 to 16 years of age. All animals were housed as continuous full-contact female-male pairs, at the Southwest National Primate Research Center (SNPRC), Texas Biomedical Research Institute, an AAALAC accredited institution. Animals were maintained under standardized marmoset husbandry conditions (Layne-Colon, Goodroe, & Burns, 2019), and received base diets (Harlan Teklad marmoset purified diet and Mazuri marmoset diet) as described in Tardif et al. (2009). The study was approved by the Texas Biomedical Research Institute Animal Care and Use Committee, and adhered to the American Society of Primatologists (ASP) Principles for the Ethical Treatment of Non-Human Primates. The sample included data from animals ranging between 333 and 629 grams of body weight. Whole-body composition was assessed with QMR, as previously described (Power et al., 2012; Tardif et al., 2009). Unsedated animals were placed in the QMR for a two-minute scan. Variables generated from the whole-body composition analysis included lean mass, fat mass, fat percentage, and fat-lean mass ratio.

### Body Condition Scoring System

We adapted and tested a body condition scoring (BCS) system for marmosets, suggested by Burns and Wachtman (2019), and based on a BCS system originally validated for rhesus macaques (Clingerman & Summers, 2005, 2012; Summers et al., 2012). The marmoset BCS system was structured around recommendations made by experienced SNPRC staff (i.e., veterinarians, veterinary technologists, colony managers and researchers), and specific to marmoset anatomy and fat deposition patterns. The procedure involves visual inspection and palpation of fat and lean tissue at thoracolumbar (scapular, spinous processes, and ribs), abdominal, and pelvic landmarks (Table 1). BCS values range from 1 (lowest) to 5 (highest), with a mid-score value of 3 representing appropriate body condition (i.e., appropriate adiposity and muscularity), and scores below or above 3 representing deleterious states of body condition. Specifically, scores below 3 represent insufficient body condition (i.e., insufficient adiposity and/or muscularity), and scores above 3 represent excessive body condition (i.e., excessive adiposity) (Table 1 and Figure 1).

**Figure 1.**
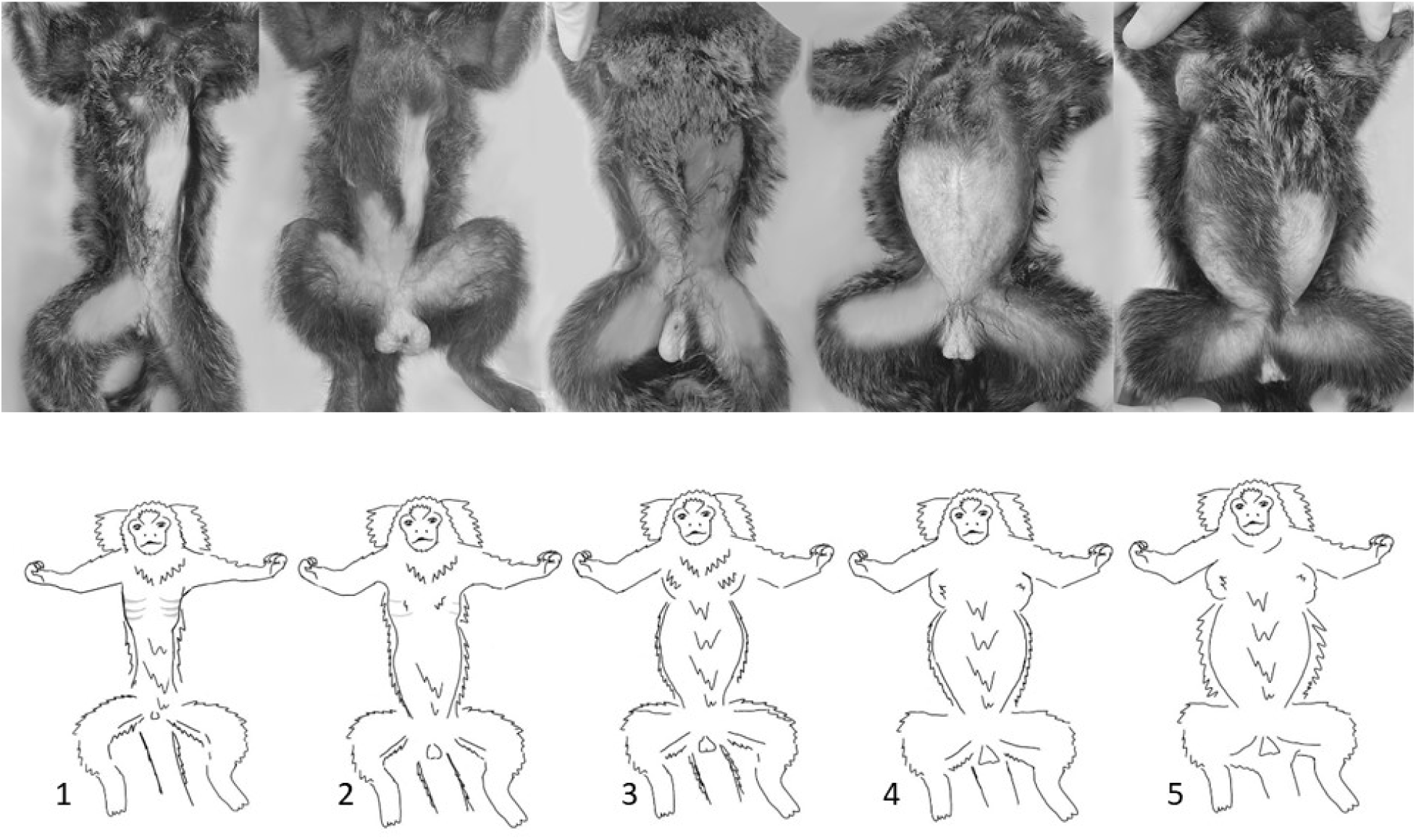
Ventral view of marmosets exhibiting body condition scores of 1 through 5 (left to right). Reference photos and drawings of BCS levels used at SNPRC are derived from the general colony population and do not include animals sampled for this study. Photos were altered to remove identifier marks (tattoos) and to remove the background substrate.

**Table 1:**
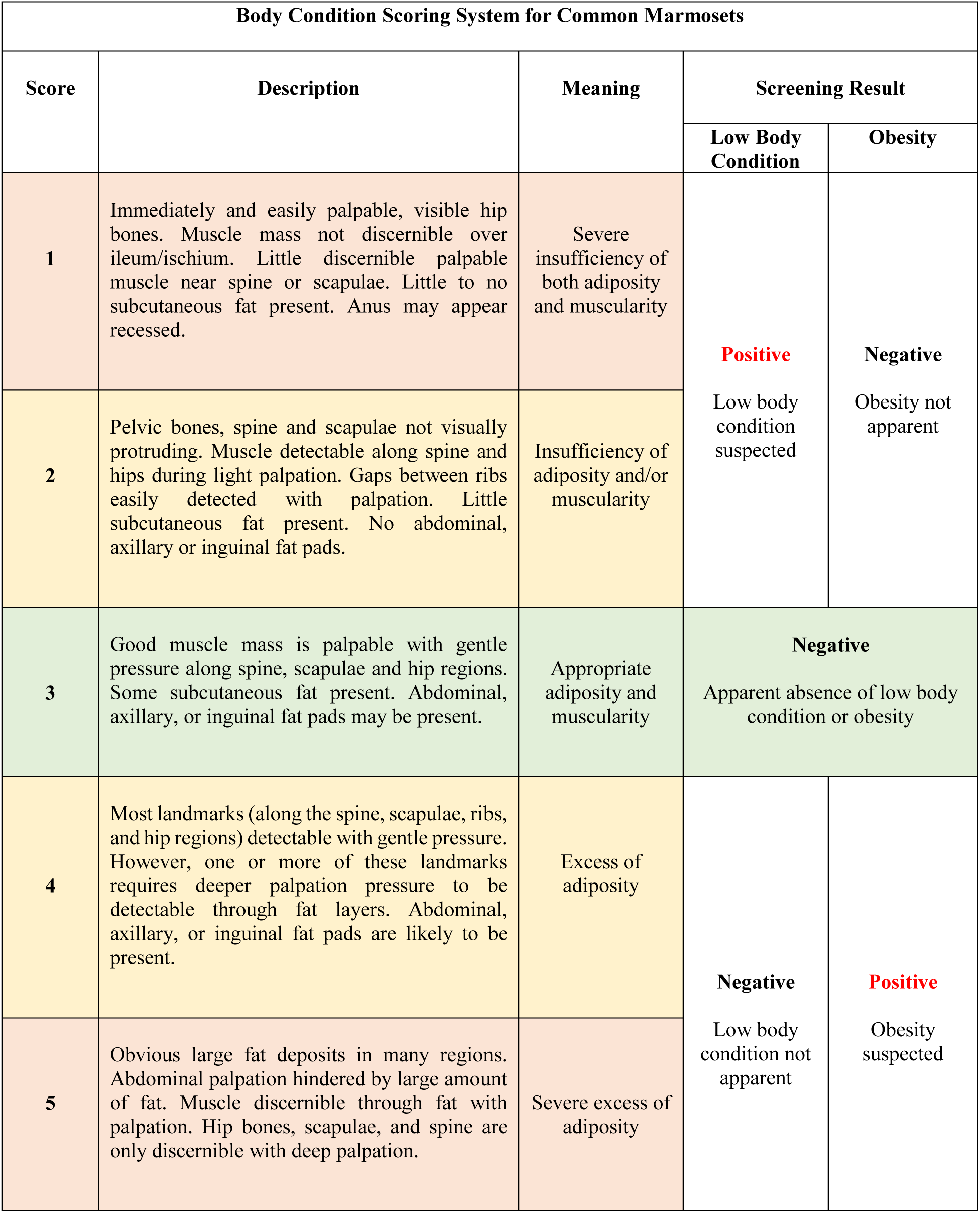
Body Condition Scoring System for Common Marmosets.

Animals were rated while unsedated, although rating can also be done under sedation at routine physical exams. The advantage of working with marmosets is that this scoring system can be performed with an unsedated animal held by a person wearing handling gloves, while a second person palpates and rates body condition (Figure 2). During data collection, raters used the descriptions shown in Table 1 to assess body condition. The score of each animal was based on the consensus of two experienced scoring staff members. Subsequently, we created a descriptive guide that provides meaning to numerical scores (Table 1). We opted for the use of neutral language and refrained from using adjectives that might introduce biases, such as “emaciated”, “thin”, “optimal” “heavy”, or “obese”; as these words might mean different things to different raters.

**Figure 2.**
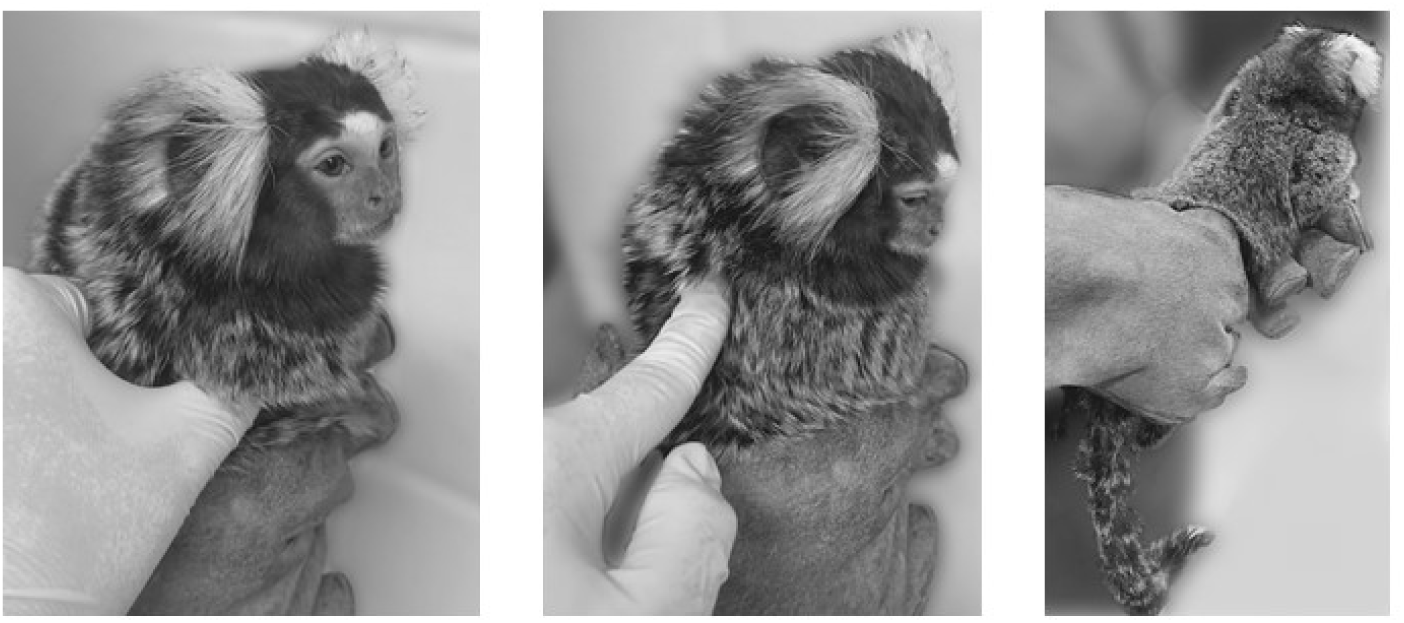
Demonstration of holding technique for alert assessment of body condition scoring of marmosets in a gloved hand to assess the ribs and spine. Photos were altered to blur the background.

### Statistical Analysis

The first objective of this study was to determine whether the marmoset BCS system accurately predicts body weight and QMR body composition parameters (i.e., lean mass, fat mass, fat percentage, and fat-lean mass ratio). First, we conducted MANOVA to test whether body weight and body composition parameters differ between BCS categories 1, 2, 3, 4, and 5. Then, we employed ordinal regression to test whether ordinal scores of BCS (i.e., body condition scores as progressive levels from 1 to 5) predict the continuous variables of body weight and body composition. Finally, we conducted binary logistic regression to test whether animals categorized as low BCS (< 3) had higher odds of exhibiting low weight and low body composition (< 1 SD below the mean), relative to animals with BCS 3 or higher. We also tested whether animals categorized as high BCS (> 3) had higher odds of exhibiting obesity (≥10% body fat), relative to animals with BCS 3 or lower.

The second objective of the study was to evaluate the performance of the marmoset BCS system as a screening test for detecting individuals exhibiting deleterious extremes of body condition: 1) insufficient adiposity and/or muscularity, or 2) excessive adiposity. This was done by making comparisons between the results of the marmoset BCS system (as a screening test), against the results produced with QMR body composition analysis (as the gold standard diagnostic test). Screening tests generate binary results, that are either positive (i.e., presence of disease) or negative (i.e., absence of disease) (Maxim et al., 2014; Trevethan, 2017). A rating of BCS 3 indicates appropriate adiposity and muscularity, and therefore represents a negative result (i.e., absence of disease) when screening for the two extreme conditions: 1) insufficient adiposity and/or muscularity, and 2) excessive adiposity. Ratings below BCS 3 indicate insufficient adiposity and/or muscularity, and consequently represent a positive result when screening for the lower extreme condition. Ratings above BCS 3 indicate excessive adiposity, and therefore represent a positive result when screening for the higher extreme condition (Table 1).

In order to evaluate BCS as a screening test for excessive adiposity, sampled animals had to be diagnosed for obesity with a gold standard method. QMR body composition analysis was used for the diagnosis and categorization of marmosets with obesity, following the criterion of 10% or higher body fat (Power et al., 2012). Likewise, assessing BCS as a screening test for low body condition required marmosets with low body condition to be diagnosed by QMR body composition analysis. However, unlike the threshold for obesity which is well established (Power et al., 2012), lower thresholds for lean and fat mass have not been established for marmosets yet. Therefore, body composition data (i.e., lean mass, fat mass, fat percentage, and fat-lean mass ratio) generated by QMR were standardized into z-scores to diagnose and categorize individuals with low body condition. Z-score values that were < 1 SD below the mean were categorized as “low”. This cutoff point was selected to represent early stages of body condition decline, as we intended to test the validity of BCS as a screening test for the presence of low body condition rather than as a diagnostic test for severe cases of insufficient adiposity and/or muscularity (Maxim et al., 2014; Trevethan, 2017)..

Screening for low body condition involved categorizing animals as either having “low body condition” or “no low body condition”. Screening for obesity involved categorizing animals as either having “obesity” or “no obesity”. The binary results (i.e., presence or absence of disease) produced by the BCS system were compared against confirmed cases of low body condition and confirmed cases of obesity, as diagnosed by QMR body composition analysis. The process of validation included screening metrics of sensitivity, specificity, positive predictive value, negative predictive value, and accuracy (Table 3) (Maxim et al., 2014; Trevethan, 2017). Data were analyzed with *R* version 4.1 (R_Core_Team, 2021) in statistical spreadsheet *jamovi* version 2.3.17 (The_jamovi_project, 2022), with *epiR: Tools for the Analysis of Epidemiological Data*. [R package] (Mark Stevenson, 2020), and *ClinicoPath jamovi Module* (Balci, 2022).

## Results

### BCS as Predictor of Body Weight and Body Composition

The sample was composed of marmosets with scores ranging from BCS 2 to BCS 5. None of the sampled animals received BCS 1. Four animals were rated as BCS 2, forty-two as BCS 3, fifteen as BCS 4, and seven as BCS 5. There were no significant differences in weight or body composition parameters between females and males (*F*(5, 62) = 0.51, *p* > 0.05). All four animals rated as BCS 2 were aged (i.e., older than 8 years) and significantly older (*F*(3, 62) = 3.83, *p* < 0.05) than animals rated as BCS 3 or higher.

Overall, the results of MANOVA indicated that ordinal levels of BCS exhibited positive relationships with body weight and all body composition variables measured by QMR. Higher BCS predicted significantly higher values in body weight and body composition (*F*(15, 166) = 7.51, Wilks’ Λ = 0.240, *p* < 0.001). In ordinal regression analyses, animals scored as BCS 2 exhibited mean values for body weight and lean mass that were below the low QMR body composition threshold, but mean values for fat parameters were above the low threshold. BCS 2 predicted lower values in body weight (*F*(3, 64) = 34.7, *p* < 0.05) and lower values in lean mass, (*F*(3, 64) = 18.9, *p* < 0.05) than BCS levels 3 through 5 (Table 2. Figures 3 and 4). BCS 2 was similar to BCS 3 in fat mass (*B* = 3.79, (95% CI, -13.11 to 20.7), *p* = 0.65) (Table 2 and Figure 5), fat percentage (*B* = 0.370, (95% CI, -3.082 to 3.82), *p* = 0.83) (Table 2), and fat-lean mass ratio, (*B* = 0.005, (95% CI, -0.04 to 0.05), *p* = 0.82) (Table 2).

**Figure 3.**
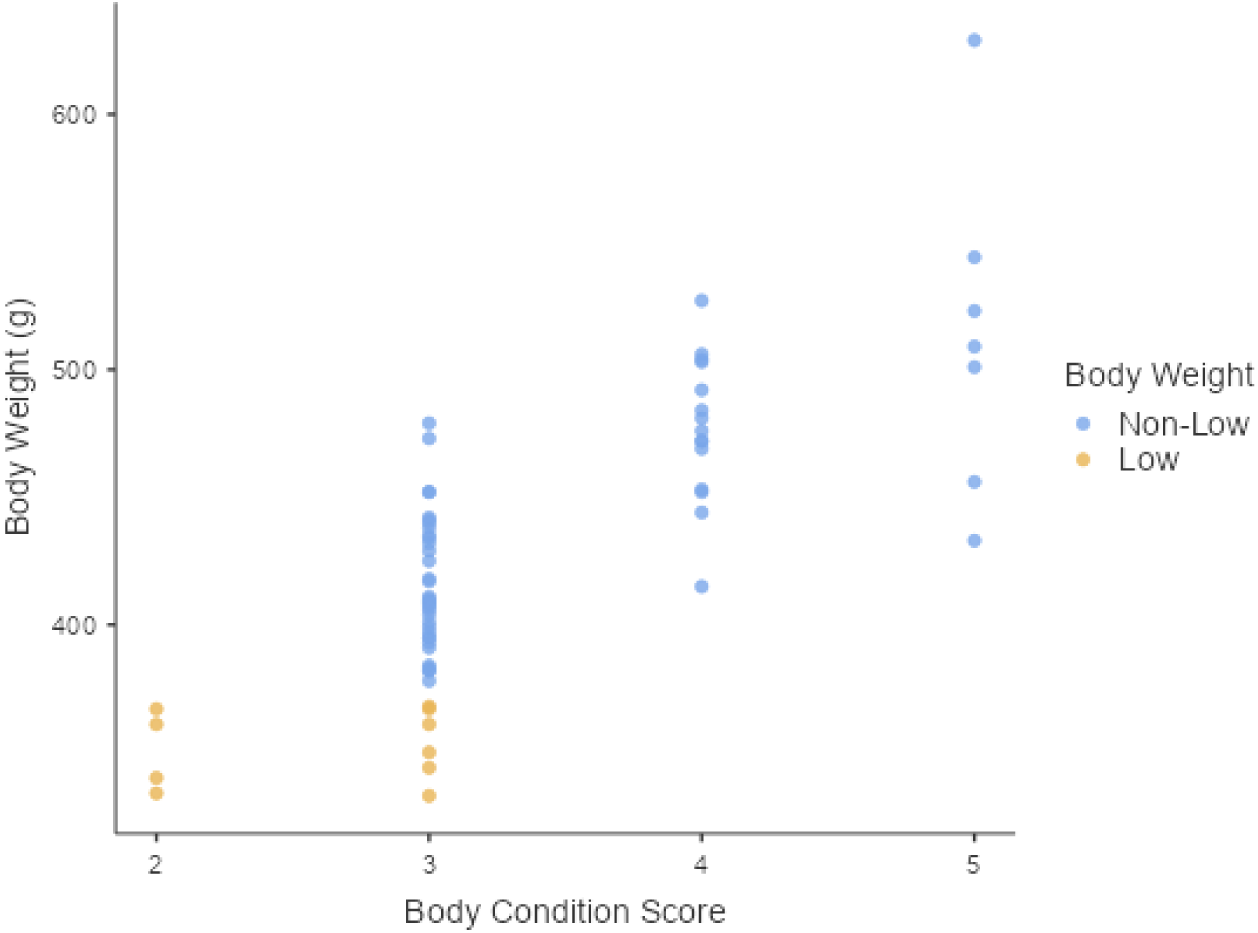
Marmoset body condition score was significantly associated with body weight. BCS 2 predicted lower values in body weight (*F*(3, 64) = 34.7, *p* < 0.05). Ordinal levels of BCS exhibited increments in body weight from BCS 2 to BCS 5. Animals with low body weight (yellow) were scored as BCS 2 or 3, and all four animals scored as BCS 2 had low body weights.

**Figure 4.**
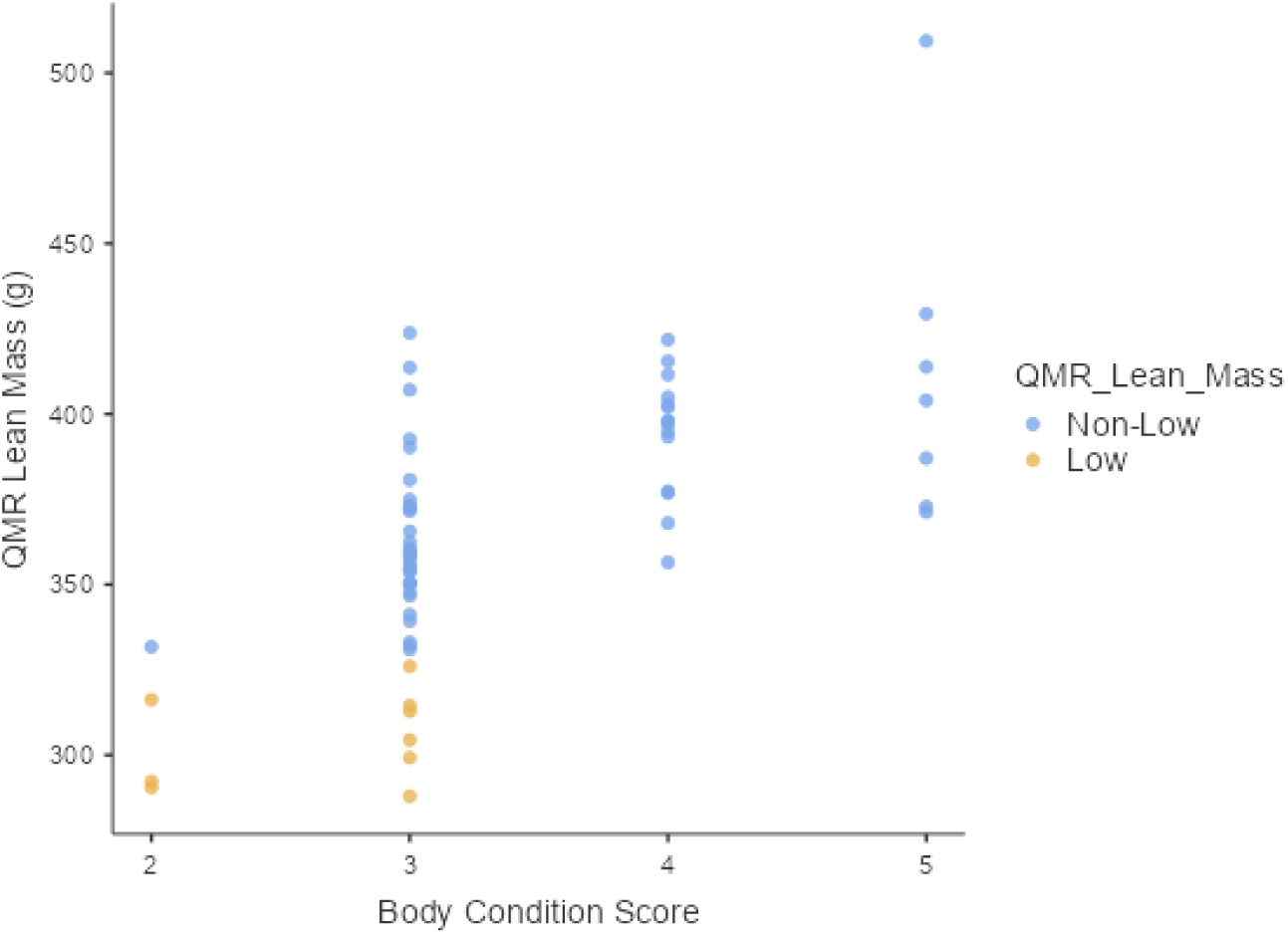
Marmoset body condition score was significantly associated with lean mass. BCS 2 predicted lower values in lean mass (*F*(3, 64) = 18.9, *p* < 0.05). Ordinal levels of BCS exhibited increments in lean mass from BCS 2 to 5. Animals with low lean mass (yellow) were scored as BCS 2 or 3, and three of the four animals scored as BCS 2 had low lean mass.

**Figure 5.**
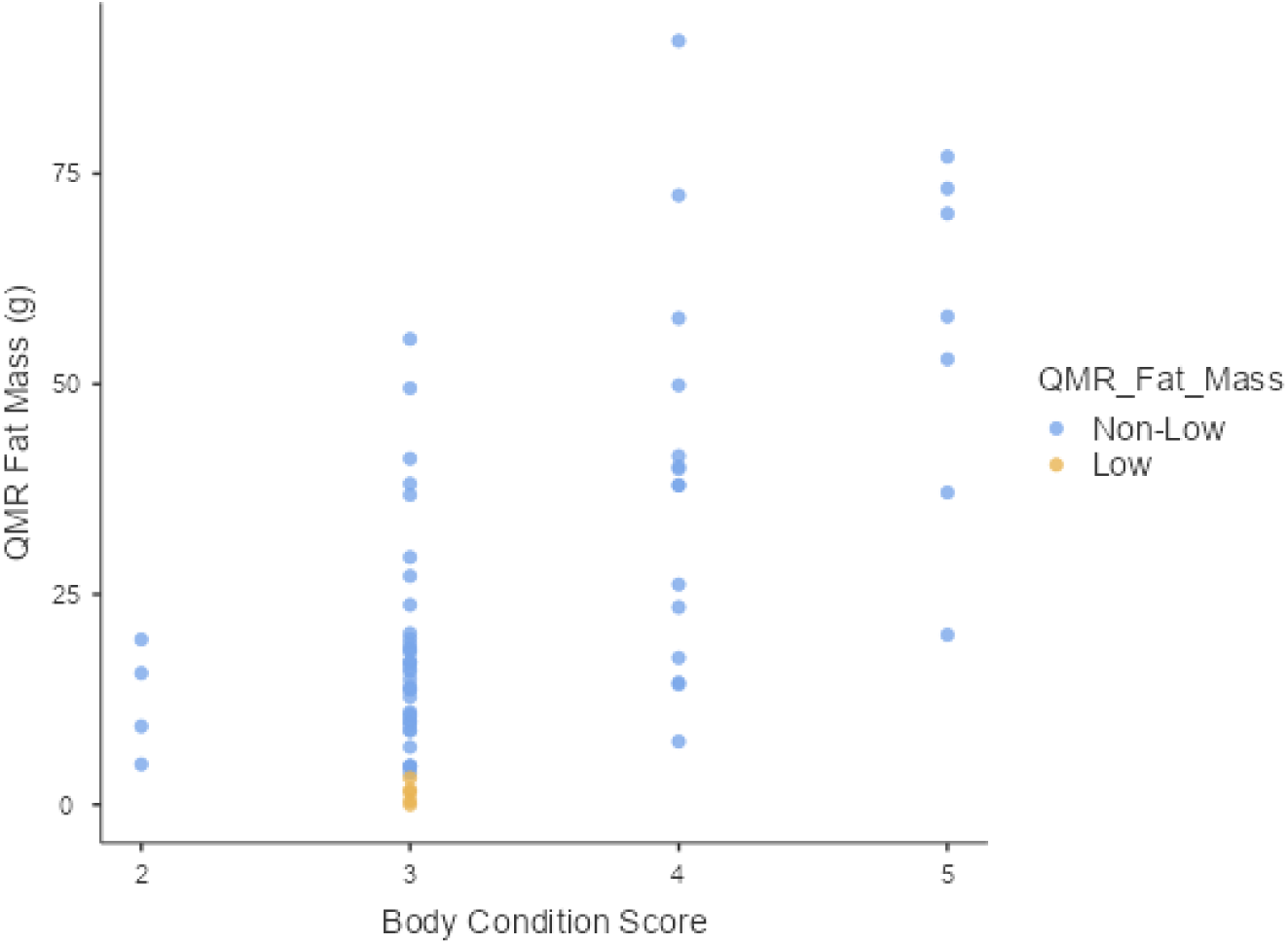
Marmoset body condition score was significantly associated with fat mass. BCS 2 predicted lower values in fat mass (*F*(3, 64) = 16.6, *p* < 0.001) than BCS 4 (*B* = 25.77, (95% CI, 7.58 to 43.9), *p* = 0.006) and BCS 5 (*B* = 43.19, (95% CI, 22.94 to 63.4), *p* < 0.001). BCS 2 was similar to BCS 3 (*B* = 3.79, 95% CI, -13.11 to 20.7, *p* = 0.66). Although ordinal levels of BCS exhibited increments in fat mass from BCS 2 to BCS 5, all animals with low fat mass (yellow) were scored as BCS 3.

**Table 2.** Summary of results for body weight and QMR body composition analysis for all marmosets, by body condition score [mean ± stdev (range)].

| Body condition score | Age (years) | Body weight (g) | Lean mass (g) | Fat mass (g) | Fat percentage | Fat-Lean mass ratio |
| --- | --- | --- | --- | --- | --- | --- |
| **2 (n=4)** | 12.75 ± 2.22**<br>(10-15) | 350.5 ± 15.97<br>(334-367) | 307.68 ± 19.89<br>(290.48-331.76) | 12.35 ± 6.59<br>(4.8-19.64) | 3.54 ± 1.87<br>(1.31-5.44) | 0.04 ± 0.02<br>(0.01-0.06) |
| **3 (n=42)** | 7.88 ± 3.08<br>(2-16) | 406.74 ± 32.5*<br>(333-479) | 354.92 ± 28.27*<br>(287.92-423.79) | 16.14 ± 12.76<br>(0-55.34) | 3.91 ± 2.95<br>(0-12.24) | 0.05 ± 0.04<br>(0-0.15) |
| **4 (n=15)** | 7.29 ± 2.87<br>(3-12) | 476.67 ± 28.39*<br>(415-527) | 394.57 ± 17.91*<br>(356.52-421.81) | 38.11 ± 22.9*<br>(7.51-90.78) | 7.83 ± 4.32*<br>(1.6-17.23) | 0.09 ± 0.06*<br>(0.02-0.23) |
| **5 (n=7)** | 7.57 ± 2.44<br>(3-10) | 513.57 ± 63.67*<br>(433-629) | 412.55 ± 47.73*<br>(371.26-509.4) | 55.54 ± 20.79*<br>(20.18-77.02) | 10.59 ± 3.38*<br>(4.66-15.13) | 0.13 ± 0.05*<br>(0.05-0.2) |

\*Statistically significant mean differences between BCS 2 and higher ordinal BCS scores.
\*\* Statistically significant mean differences in age between BCS scores.

For fat mass, BCS 2 (*F*(3, 64) = 16.6, *p* < 0.001) predicted lower values in comparison to BCS 4 (*B* = 25.77, (95% CI, 7.58 to 43.9), *p* = 0.006) and BCS 5 (*B* = 43.19, (95% CI, 22.94 to 63.4), *p* < 0.001) (Table 2 and Figure 5). For fat percentage, BCS 2 (*F*(3, 64) = 11.7, *p* < 0.001), predicted lower values in comparison to BCS 4 (*B* = 4.283, (95% CI, 0.571 to 8.00), *p* = 0.02) and BCS 5 (*B* = 7.05, (95% CI to 2.92, 11.18), *p* = 0.001) (Table 2). BCS 2 predicted lower values of fat-lean mass ratio (*F*(3, 64) = 12.2, *p* < 0.001) in comparison to BCS 4 (*B* = 0.06, (95% CI, 0.01 to 0.103), *p* = 0.021) and BCS 5 (*B* = 0.092, (95% CI, 0.03 to 0.14), *p* < 0.001) (Table 2). Adjusted percentages of variance explained by BCS were 60.1% in body weight, 44.5% in lean mass, 41.2% in fat mass, 32.4% in fat percentage, and 33.4% in fat-lean mass ratio.

Binary logistic regression indicated that low BCS (i.e., BCS 2) significantly predicted low lean mass (χ^2^ (1) = 8.83, *p* = 0.003). Animals scored as BCS 2 had higher odds of exhibiting low lean mass (*B* = 3.37, (95% CI, 0.95 to 5.78), OR = 29.0, *p* = 0.006), relative to animals with BCS of 3 or higher. However, BCS 2 did not predict low body weight, low fat mass, low fat percentage, or low fat-lean mass ratio (*p* > 0.05). In the assessment of high BCS (i.e., BCS 4 or BCS 5) as a predictor of obesity (i.e., ≥10% body fat) with binary logistic regression, high BCS predicted obesity status (χ^2^ (1) = 14.0, *p* < 0.001). Animals that scored BCS 4 or 5 had higher odds of having obesity (*B* = 2.72, (95% CI, 1.07 to 4.38), OR = 15.23, *p* = 0.001), relative to animals scored as BCS 2 or 3 (Table 3).

**Table 3.** Obesity (≥10% body fat) status by body condition score.

| Body condition score | QMR <10% body fat | QMR $\geq 10\%$ body fat |
| --- | --- | --- |
| 2 (n = 4) | 4 | 0 |
| 3 (n = 42) | 40 | 2 |
| 4 (n = 15) | 11 | 4 |
| 5 (n = 7) | 2 | 5 |

### Accuracy

In comparison to the gold standard QMR body composition analysis, the accuracy of the BCS system as a screening test for low body condition ranged between 79.4% and 91.2%.

### Sensitivity

The BCS test had low sensitivity. Individuals with low body condition as measured by QMR, have a decreased probability of being detected as “positives” for this condition by the BCS screening test alone. Among individuals with low body condition, the 95% confidence interval probabilities of being detected (i.e., obtaining a positive BCS screen test result) were 12.2% - 73.8% for low body weight, 7.5% - 70.1% for low lean mass, 0% - 52.2% for fat mass, 0.3% - 44.5% for low fat percentage, 0% - 33.6% for fat-lean mass ratio, and 6.4% - 47.6% overall.

### Specificity

The BCS test showed high specificity. Individuals with non-low body condition as measured by QMR, have an increased probability of being rated as above 2 in the BCS screening test. Among individuals with non-low body condition, the 95% confidence interval probabilities of being correctly identified as “negative” for low body condition by the BCS screening test were 93.8% - 100% for body weight, 90.9% - 100% for lean mass, 84.5% - 98.2% for fat mass, 85.6% - 98.9% for fat percentage, 83.5% - 98.1% for fat-lean mass ratio, and 92.9% - 100% overall.

### Positive Predictive Value (PPV)

PPV varied by parameter. Among individuals with a positive BCS screening test result for low body condition, the probabilities of being a true positive case were 100% for low body weight, 75% for low lean mass, 0% for low fat mass, 25% for low fat percentage, 0% for low fat-lean mass ratio, and 100% overall.

### Negative Predictive Value (NPV)

The test showed high NPV. Among individuals with a negative BCS screening test result for low body condition, the probabilities of being a true negative were 90.6% for body weight, 90.6% for lean mass, 92.2% for fat mass, 85.9% for fat percentage, 85.9% for fat-lean mass ratio, and 78.1% overall.

**Table 4.**
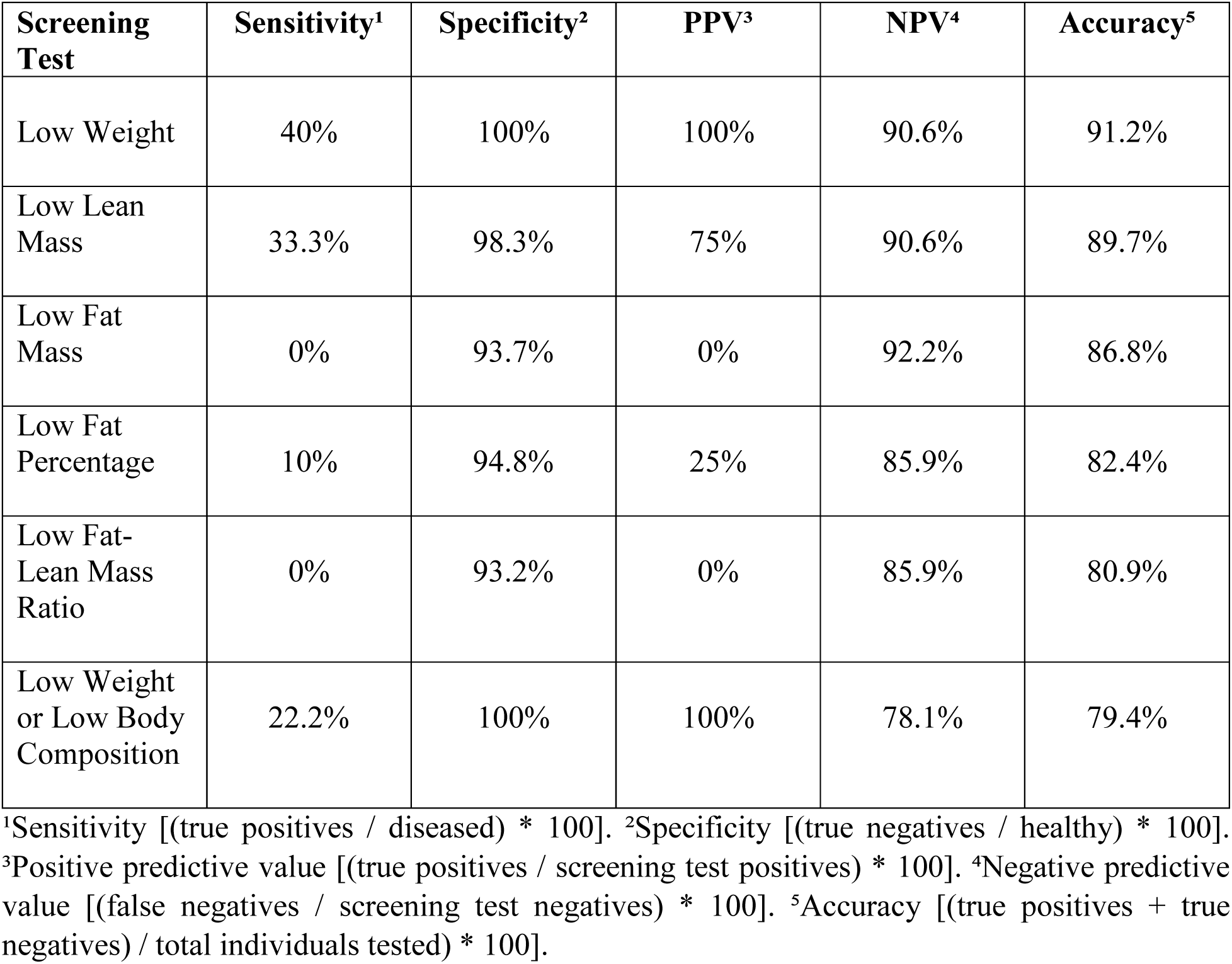
BCS as a screening test for low body condition (< 1 SD below mean)

| **Screening Test** | **Sensitivity<sup>1</sup>** | **Specificity<sup>2</sup>** | **PPV<sup>3</sup>** | **NPV<sup>4</sup>** | **Accuracy<sup>5</sup>** |
| --- | --- | --- | --- | --- | --- |
| Low Weight | 40% | 100% | 100% | 90.6% | 91.2% |
| Low Lean Mass | 33.3% | 98.3% | 75% | 90.6% | 89.7% |
| Low Fat Mass | 0% | 93.7% | 0% | 92.2% | 86.8% |
| Low Fat Percentage | 10% | 94.8% | 25% | 85.9% | 82.4% |
| Low Fat-Lean Mass Ratio | 0% | 93.2% | 0% | 85.9% | 80.9% |
| Low Weight or Low Body Composition | 22.2% | 100% | 100% | 78.1% | 79.4% |

<sup>1</sup>Sensitivity [(true positives / diseased) \* 100]. <sup>2</sup>Specificity [(true negatives / healthy) \* 100].
<sup>3</sup>Positive predictive value [(true positives / screening test positives) \* 100]. <sup>4</sup>Negative predictive value [(false negatives / screening test negatives) \* 100]. <sup>5</sup>Accuracy [(true positives + true negatives) / total individuals tested) \* 100].

The prevalence of obesity (≥10% body fat) as assessed by QMR in this sample was 16.2% (11/68). In comparison to QMR fat percentage, the accuracy of BCS as screening test for obesity was 77.9%. The screening test for obesity exhibited a sensitivity of 81.8%. Animals with obesity have a high probability of being detected as “positives” by the BCS screening test, as nine out of eleven cases of obesity were correctly identified. Among individuals with obesity, the 95% confidence interval probabilities of being detected were 48.2% - 97.7%.

The screening test for obesity had a specificity of 77.2%. Animals without obesity have an acceptable to good probability of obtaining a negative result in the BCS screening test, with forty-four out of fifty-seven animals without obesity being correctly identified. Among animals without obesity, the 95% confidence interval probabilities of being correctly identified as “negative” by the BCS screening test were 64.2% - 87.3%.

The obesity screening test had a low PPV. Among animals with a positive BCS screening test result, the probability of being a true positive case for obesity was 40.9%. Nine out of twenty-two positives were true positives. The screening test for obesity exhibited a high NPV. Among animals with a negative BCS screening test result, the probabilities of being a true negative (i.e., not having obesity) were 95.7%. Forty-four out of forty-six negatives were true negatives.

**Table 5.** BCS as a screening test for obesity (≥10% body fat)

| **Screening Test** | **Sensitivity<sup>1</sup>** | **Specificity<sup>2</sup>** | **PPV<sup>3</sup>** | **NPV<sup>4</sup>** | **Accuracy<sup>5</sup>** |
| --- | --- | --- | --- | --- | --- |
| Obesity | 81.8% | 77.2% | 40.9% | 95.7% | 77.9% |

<sup>1</sup>Sensitivity [(true positives / diseased) \* 100]. <sup>2</sup>Specificity [(true negatives / healthy) \* 100].
<sup>3</sup>Positive predictive value [(true positives / screening test positives) \* 100]. <sup>4</sup>Negative predictive value [(false negatives / screening test negatives) \* 100]. <sup>5</sup>Accuracy [(true positives + true negatives) / total individuals tested) \* 100].

## Discussion

### BCS as predictor of body weight and body composition

We aimed to evaluate whether our suggested marmoset BCS system predicts body weight and body composition, and to assess its performance as a screening test for low body condition and obesity in marmosets. Overall, BCS is predictive of body weight and body composition parameters, suggesting that it can be used as a screening test for detecting low body condition and obesity in marmosets. For this assessment, we sampled females and males between 2 and 16 years of age. We did not find sexual dimorphism in weight or whole-body composition parameters. This finding suggests that the same thresholds and categories can be employed to assess both female and male marmosets. Our findings also suggest that the BCS system is a useful tool for monitoring aging marmosets. Animals rated as having low body condition (BCS 2) were significantly older than animals rated as BCS 3 or higher, highlighting the aging-related decline in body condition, and the importance of screening geriatric marmosets.

The BCS system predicted incremental differences in body weight and lean mass across ordinal BCS levels 2 through 5. Therefore, BCS allows raters to predict an animal’s body condition status in terms of body weight and lean mass. Although this BCS system predicted differences in adiposity (i.e., fat mass, fat percentage, and fat-lean mass ratio) between BCS 2 and BCS 4 to 5, it did not predict differences between BCS 2 and BCS 3. This finding suggests that in terms of palpating adiposity, human fingers are not sensitive enough to accurately differentiate between animals with insufficient body condition (BCS 2) and animals with appropriate body condition (BCS 3). Animals scored as BCS 2, had higher odds of exhibiting low lean mass, but not low body weight or low adiposity, relative to animals with BCS 3 or higher. Therefore, the results strongly suggest that this BCS system is significantly better at predicting low lean mass, than at predicting low adiposity in marmosets.

As the BCS system predicted incremental differences in adiposity at the higher levels of body condition (BCS 4 and 5), it was also a predictor for excess of adiposity. Animals with BCS 4 and BCS 5, exhibited higher odds of obesity based on the ≥10% body fat threshold (Power et al., 2012). This finding suggests that although human fingers might not be sensitive enough to differentiate adiposity between BCS 2 and BCS 3, sensitivity improves at higher levels of adiposity, allowing raters to accurately differentiate between animals with BCS 3 and BCS 4 to 5. Therefore, our results suggest that the BCS system is better at predicting excessive adiposity, than at predicting insufficient adiposity in marmosets.

### BCS as a screening test for low body condition

The BCS screening test was slightly better at detecting cases of low body weight and low lean mass, than detecting cases of low adiposity (i.e., low fat mass, low fat percentage, and low fat-lean mass ratio). Nonetheless, the BCS screening test for low body condition has high specificity, and produced a high number of true negatives. Therefore, a positive result provides sufficient confidence to “rule in” the presence of low body condition.

### BCS as a screening test for obesity

The BCS screening test for obesity is sensitive enough to detect most of the true positive cases of obesity, and produced a low number of false negatives. A negative result (BCS below 4) might be sufficient to “rule out” the presence of obesity in most instances, as less than 5% of marmosets with a score of 3 had percent body fat above 10%, and none with a score of 2. The BCS screening test for obesity has acceptable specificity, and produced a high number of true negatives. Based on this, a positive result provides acceptable but limited confidence to “rule in” the presence of obesity. The screening test is good at correctly identifying animals without obesity. False negatives occurred at low frequencies, and individuals with a negative result are highly likely to be true negatives.

### BCS as predictor and screening test for low body condition and obesity

After an examination of the relationships between ordinal levels of BCS and each body composition variable, the results suggest that it is primarily lean mass, and not fat mass, that allows the BCS system to serve as an accurate screening test for detecting animals with low body condition. We have noticed that when palpating an animal, muscle tissue typically feels denser or more substantial than subcutaneous adipose tissue. It seems that during palpation, the presence (tactile sensation) of muscle mass contributes at a higher degree than the presence (tactile sensation) of adipose tissue, to the rater’s sensorial perception of body condition. Thus, it seems that when we palpate over skeletal protuberances, we are relying more on feeling muscle tissue than feeling fat tissue to score animals. It is plausible that muscle tissue hides skeletal protuberances better than subcutaneous adipose tissue, and therefore palpation through muscle might require the rater to apply higher pressure with the fingers to reach bone. In contrast, palpation through subcutaneous adipose tissue might present less resistance to the rater, allowing fingers to sink in and feel bone more easily.

Although BCS is useful as a screening tool for detecting low body condition and obesity in marmosets, it has limitations. The descriptions in this BCS system instruct the rater to observe and palpate muscle and fat. However, the abundance of muscle and fat is assessed in a single score, making no distinction between them, rather than assigning separate scores for muscle and fat. This represents a limitation, in comparison to body composition analysis. We suggest it might be possible to create a BCS system with higher accuracy, by having two separate scores, one for muscularity and one for adiposity. Another potential limitation could arise if BCS produces different results when an animal is sedated vs. unsedated. If differences in muscle tension between sedated and unsedated states can alter the tactile perception of raters, then this may lead to scoring discrepancies. All animals in this study were rated while unsedated, and therefore we did not test for agreement between sedated and unsedated ratings. More research is required to determine if sedation or separation of scores (muscularity and adiposity) improves accuracy as a screening tool.

For this assessment, we defined “low body condition” as values in body weight and values in body composition that were < 1 SD below the mean. Although it might be possible to increase the sensitivity of the BCS screening test by setting the cutoff point of low body condition to values < 2 SD below the mean, our sample included few cases with values < 2 SD below mean. Therefore, we did not assess the validity of the BCS test in marmosets with severely low body condition. Instead, we evaluated a sample of relatively healthy animals, with individuals exhibiting mild to moderate reductions in body condition. This is an important distinction, because the usefulness of BCS as a screening test for low body condition, depends on its ability to accurately identify individuals at early stages of body condition decline.

This assessment shows that the BCS system for marmosets is significantly better at predicting low lean mass than at predicting low adiposity, and significantly better at predicting obesity than at predicting low adiposity. At lower levels of adiposity, the BCS system appears to allow raters to detect animals with low body condition more accurately by relying on the palpation of lean mass rather than adipose tissue. At higher levels of adiposity, the BCS system allows raters to accurately identify animals with obesity. Full validation of BCS as a screening test for low body condition and obesity in common marmosets will rely on future assessments of intra-rater consistency and inter-rater agreement (intra and inter-rater reliability).

Overall, the BCS screening test exhibits acceptable to good accuracy in comparison to the gold standard of QMR body composition. The BCS test can serve as a low cost, quick, non-invasive, instrument-free method to screen for low body condition and obesity in common marmosets. It is important to highlight that the BCS test is a subjective semi-quantitative screening test, not a diagnostic test (Maxim et al., 2014; Trevethan, 2017), and as such it cannot be considered a substitute for objective measurements of body weight, body composition, or clinical diagnostic tests. The BCS screening test can be used to monitor changes in body condition and to alert caretakers, clinicians, and researchers about possible changes in health. Suspicion of illness, wasting, or excessive adiposity can be followed up with diagnostic testing.

## Abbreviations

BCS: body condition scoring
QMR: quantitative magnetic resonance
PPV: positive predictive value
NPV: negative predictive value

## Author Contributions

**Juan Pablo Arroyo:** conceptualization (equal); data curation (lead); formal analysis (lead); investigation (lead); methodology (lead); project administration (supporting); supervision (supporting); validation (lead); visualization (lead); writing – original draft (lead); writing – review and editing (equal).

**Addaline Alvarez:** data curation (equal); investigation (equal); methodology (equal); project administration (equal); supervision (supporting); visualization (equal); writing – review and editing (supporting).

**Lori Alvarez:** data curation (equal); investigation (equal); methodology (equal); project administration (equal); supervision (equal); visualization (supporting); writing – review and editing (supporting).

**Alexana J. Hickmott:** data curation (equal); formal analysis (supporting); investigation (equal); methodology (equal); project administration (supporting); supervision (equal); visualization (equal); writing – review and editing (supporting).

**Aaryn C. Mustoe:** formal analysis (supporting); investigation (supporting); visualization (equal); writing – review and editing (equal).

**Kathy Brasky:** conceptualization (supporting); methodology (equal); project administration (equal); supervision (equal); writing – review and editing (equal).

**Kelly R. Reveles:** conceptualization (supporting); formal analysis (supporting); funding acquisition (equal); methodology (equal); project administration (equal); supervision (equal); visualization (equal); writing – review and editing (equal).

**Benjamin J. Ridenhour:** conceptualization (supporting); formal analysis (supporting); funding acquisition (equal); methodology (equal); project administration (equal); supervision (equal); visualization (equal); writing – review and editing (equal).

**Katherine R. Amato:** conceptualization (supporting); formal analysis (supporting); funding acquisition (equal); methodology (equal); project administration (equal); supervision (equal); visualization (equal); writing – review and editing (equal).

**Michael L. Power:** conceptualization (equal); data curation (equal); formal analysis (equal); funding acquisition (equal); methodology (equal); project administration (equal); supervision (equal); validation (equal); visualization (equal); writing – review and editing (equal).

**Corinna N. Ross:** conceptualization (lead); data curation (equal); formal analysis (equal); funding acquisition (lead); methodology (equal); project administration (lead); supervision (lead); validation (equal); visualization (equal); writing – original draft (equal); writing – review and editing (lead).

## Acknowledgments

We thank Donna Lane-Colón and the marmoset care team at SNPRC. Research was approved by the Southwest National Primate Research Center Animal Care and Use Committee (IACUC #1743 CJ). This investigation used resources that were supported by NIH grants R01AG065546 and U34AG068482, and the base grant to SNPRC (P51 OD011133) from the Office of Research Infrastructure Programs, NIH.

## Ethics Statement

The authors have nothing to report.

## Data Availability statement

The data that support the findings of this study are available from the authors upon reasonable request.

